# Deconvoluted methylation profiles discriminate between closely related melanocytic nevi

**DOI:** 10.1101/2024.06.11.598516

**Authors:** Daniel Aldea, Nicolas Macagno, Elise Marechal, Mathias Moreno, Pauline Romanet, Morgane Pertuit, Jérémy Garcia, Nathalie Degardin, Stéphanie Mallet, Isabelle James, Guillaume Captier, Anne Barlier, Heather C. Etchevers

## Abstract

Congenital melanocytic nevi (CMN) and common acquired melanocytic nevi (AMN) are melanoma-predisposing skin conditions presenting excessive numbers of melanocytes but arising at different times in life. Appropriately weighted whole-genome methylation analysis can be used as a basis for further research in dermatopathology and as applied here provides new insights into nevus biology.

## TO THE EDITOR

Common or “acquired” (AMN) and congenital melanocytic nevi (CMN) are discrete accumulations of skin melanocytes distinguished by their visibility and size at birth. Both are benign but exhibit differences in presumed origin, phenotype, and implications for human health (Maher et al. 2023). AMN emerge as small (<6 mm) postnatal lesions, increasing in number during adolescence to stabilize again in adulthood. In contrast, CMN arise during fetal development. They vary in size, inversely proportional to incidence, from frequent and small (less than 1.5 cm diameter) to rare and giant (exceeding 40 cm). CMN are also diverse in terms of coarse terminal hair, texture, pigmentation dynamics, or nodules, not typical of AMN.

Malignant transformation of any nevus is rare; however, a quarter of all superficial spreading melanomas (SSM) originate from a pre-existing AMN (Pampena et al. 2017). In contrast, multiple medium to giant CMN are associated with significantly increased risk for prepubertal melanoma, particularly nodular (NM) but also extracutaneous, in direct proportion to the size and number of congenital lesions. While many studies focus on transformation of common nevi to melanoma, less is known about the molecular specificities of CMN and how these impact melanoma risk (Bastian et al. 2002).

AMN and CMN are associated with mosaicism for activating mutations at successive levels of the RAS-Mitogen activated protein kinase (MAPK) signaling pathway, which foster melanomagenesis. The second most commonly identified somatic mutation in large and giant (L/G) CMN after *NRAS* p.Q61(K/R/L) (Dessars et al. 2009), *BRAF* p.V600E predominates in small CMN and AMN (Muse et al. 2022; Zalaudek et al. 2011). CMN-associated proliferative nodules or melanoma differ in histone H3K27 trimethylation (Busam et al. 2017; Macagno et al. 2018) and propensity to chromosomal breaks (Bastian et al. 2002). While direct epigenetic marks like DNA methylation may reflect differential nevus potential by exerting stable effects on transcriptional regulation, they had not been studied in confirmed CMN.

We therefore isolated DNA from the CMN of twelve patients scheduled for excision after written, informed consent from their parents (details are in the Supplementary Material and Methods and Supplementary Table S1). Six harbored *NRAS* missense mutations at p.Q61(K or R), two at p.G13(R), and four in *BRAF* at p.V600(E). DNA methylation status was assessed using Infinium MethylationEPIC microarrays and integrated with data from a technically comparable study matching sixteen AMN with adjacent, unaffected control skin samples (Muse et al. 2022). These were from a subset of eleven participants under age 50, presenting BRAF p.V600(E, K, or R) or NRAS p.G12(R) mutations. In total, 44 samples were pooled for further analyses (computational details are in Supplementary Material and Methods and Fig. S1a). Their clinical and molecular features are compared in Supplementary Table S1.

To account for technical, clinical, and cellular variability, we first controlled for potential batch effects introduced by the different Illumina array generations. We then performed reference-based deconvolution of each methylome by predicting the proportions of melanocytes, epidermal keratinocytes, and fibroblasts relative to cell-specific methylation profiles (Muse et al. 2022). After normalization, 698,939 5′-C-phosphate-G-3′ (CpG) probes shared a cross-sample bimodal density distribution of beta values (Figure 1a; Supplementary Material and Methods and Fig. S1a). Predicted melanocyte proportions in CMN were 53.2% (S.D. ±19.1%), compared to 21.8% (S.D. ±14.7%) in AMN and 7.7% (S.D. ±2.7%) in control skin samples (Figure 1b and Supplementary Fig. S1b). We next measured true proportions of SOX10^+^ melanocytes and P63^+^ keratinocytes in typical CMN (patients 8, 9, 11 and 12) sections adjacent to those used for DNA extraction, quantifying positive immunohistochemistry relative to the pan-cellular expression of SMARCB1^+^ (INI1) in 93153, 194044, 72000 and 89530 nuclei, respectively (Figure 1c-d, Supplementary Tables S1-2; Supplementary Fig. S1c). Enumerated by machine learning in an adaptable workflow (see Supplementary Materials and Methods), these unbiased measurements validated our methylome-based predictions (Pearson correlation coefficient > 0.98) (Supplementary Fig. S1c).

**Figure 1.**
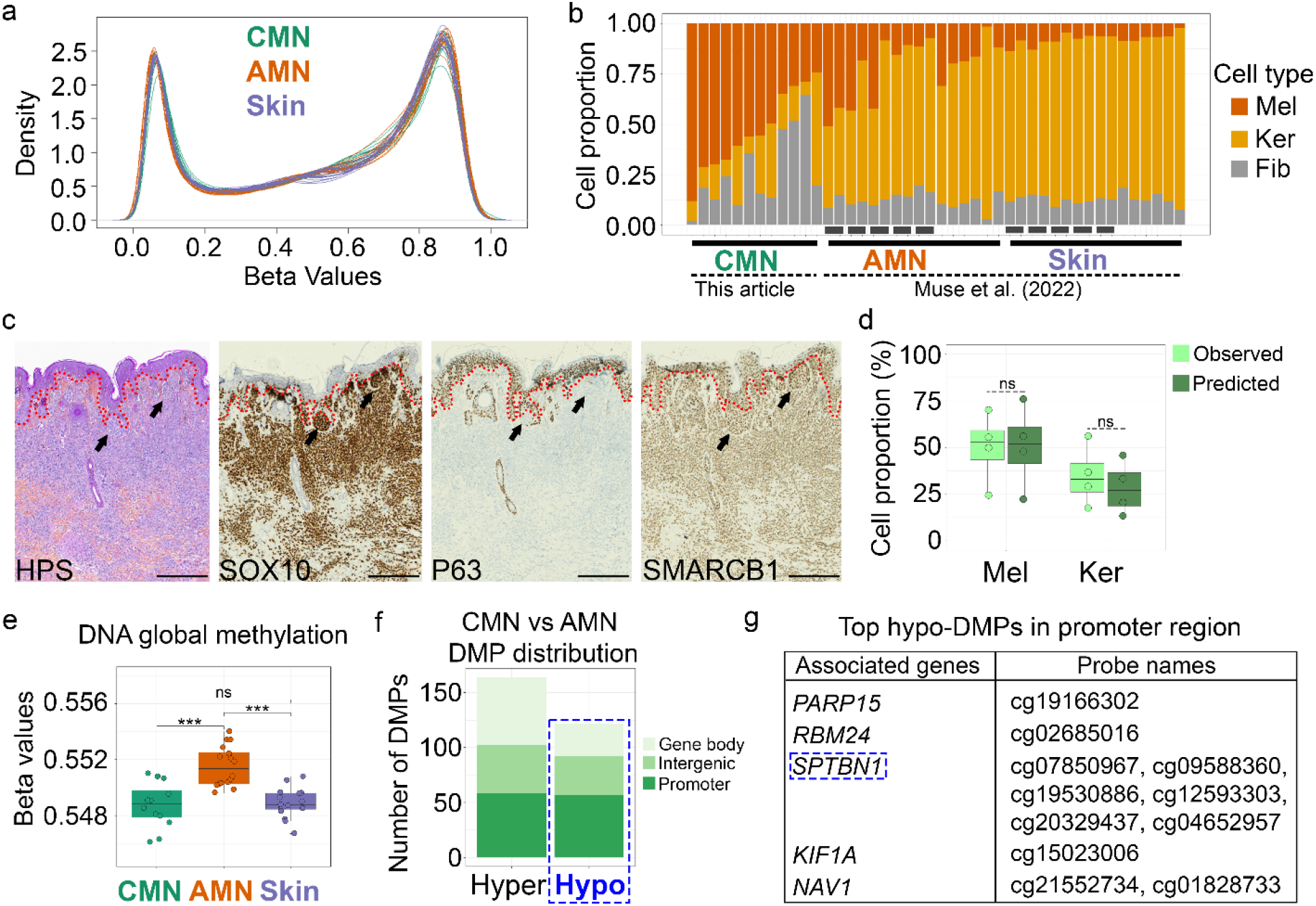
Inferred cellular architecture of congenital melanocytic nevi and epigenetic methylation in CMN vs. AMN. **(a)** Density plots of quantile-normalized and cell-deconvoluted beta values from CMN, AMN, and control Skin groups. **(b)** Estimated cellular proportions in congenital or acquired melanocytic nevi (CMN and AMN), versus control skin (Skin) for melanocytes (Mel), keratinocytes (Ker), and fibroblasts (Fib). Each vertical line represents one sample (N=44 samples). For AMN and Skin, samples from the same patient are underlined. AMN and unaffected Skin data were obtained from the Gene Expression Omnibus accession GSE188593 (Muse et al. 2022). **(c)** Representative CMN images. Hematoxylin-phloxine-saffron (HPS) staining shows high nuclear density (purple) underlying the epidermis. Immunohistochemistry staining of SOX10 for (nevo)melanocytes, P63 for epidermis and epidermal annexes, and SMARCB1 for all normal nuclei. Dotted lines delimit the epidermal-dermal junction. Melanocytic nests (black arrow) are indicated. **(d)** Boxplot showing observed *versus* predicted cellular proportions of SOX10^+^ melanocytes and P63^+^ keratinocytes in CMN, N=4 independent samples (patients 8, 9, 11, and 12). **(e)** Box plots representing global DNA methylation of CMN, AMN, and unaffected Skin. The mean of beta values across 698,939 CpG probes common to all samples is plotted. **(f)** The genomic distribution of the hyper- and hypo-DMPs is classified into 3 genomic locations; gene body, intergenic, and promoter (1500-TSS, 200-TSS, 5’UTR). **(g)** Top hypo-DMPs in CMN vs AMN within the promoter region. Significance was assessed by **(d)** a two-tailed Mann-Whitney T-test and **(e)** Kruskal-Wallis one-way ANOVA. ns. not significant, ***P<0.001.

The selected AMN methylomes (see details in Supplementary Table S1), are globally hyper-methylated relative to adjacent control skin (Muse et al. 2022), a reported finding that we confirmed after deconvolution. In contrast, deconvoluted CMN profiles were hypo-methylated compared to AMN, but not significantly different from unaffected skin (Figure 1e). Pairwise CMN, AMN, and control skin group comparisons pinpointed 286 significant differentially methylated positions (DMPs) with a False Discovery Rate (FDR) <0.05 and size effect |≥0.5| (Supplementary Materials and Methods and Table S3). Nearly two-thirds of the DMPs between CMN and control skin were located in intergenic and promoter genomic regions (Figure 1f). Differential methylation of regulatory regions may thus distinguish CMN from AMN or unaffected skin.

Among top significantly hypo-methylated CpGs in promoters between CMN and AMN (Figure 1g), the *SPTBN1* gene had six hypo-DMPs (cg07850967, cg09588360, cg19530886, cg12593303, cg20329437 and cg04652957) close to each other and an alternative first exon (Figure 2a-c, Supplementary Fig. S1e, and Supplementary Table S3). Spectrin Beta, Non-Erythrocytic 1 (SPTBN1) encodes a key subunit of a tetrameric cytoskeletal protein with homology to α-actinin, involved in cell cycle progression, differentiation, tumor suppression, and epithelial maintenance in diverse models (Yang et al. 2021). In published skin single-cell RNA-seq atlases (Solé-Boldo et al. 2020; Tirosh et al. 2016) *SPTBN1* is expressed in both normal and malignant melanocytes (Supplementary Fig. S1f). This observation led us to further investigate its protein expression in melanocytic nevi. Intense SPTBN1 immunofluorescence labeled spinous and granular epidermis of all skin samples (Figure 2d-e and Supplementary Fig. S2a-b). SPTBN1 also co-localized with SOX10^*+*^ in basal epidermal melanocytes (Figure 2d-e), further supporting our analysis from single-cell RNA-seq skin atlases, where over 9% of normal melanocytes co-express *SPTBN1* and *SOX10* (Supplementary Fig. S1f). Additionally, SPTBN1 was strongly expressed in SOX10^+^ dermal melanocytes of CMN, compared to weaker fibroblast or junctional melanocyte staining in AMN or unaffected skin (Figure 2d-e and Supplementary Fig. S2a-b).

**Figure 2.**
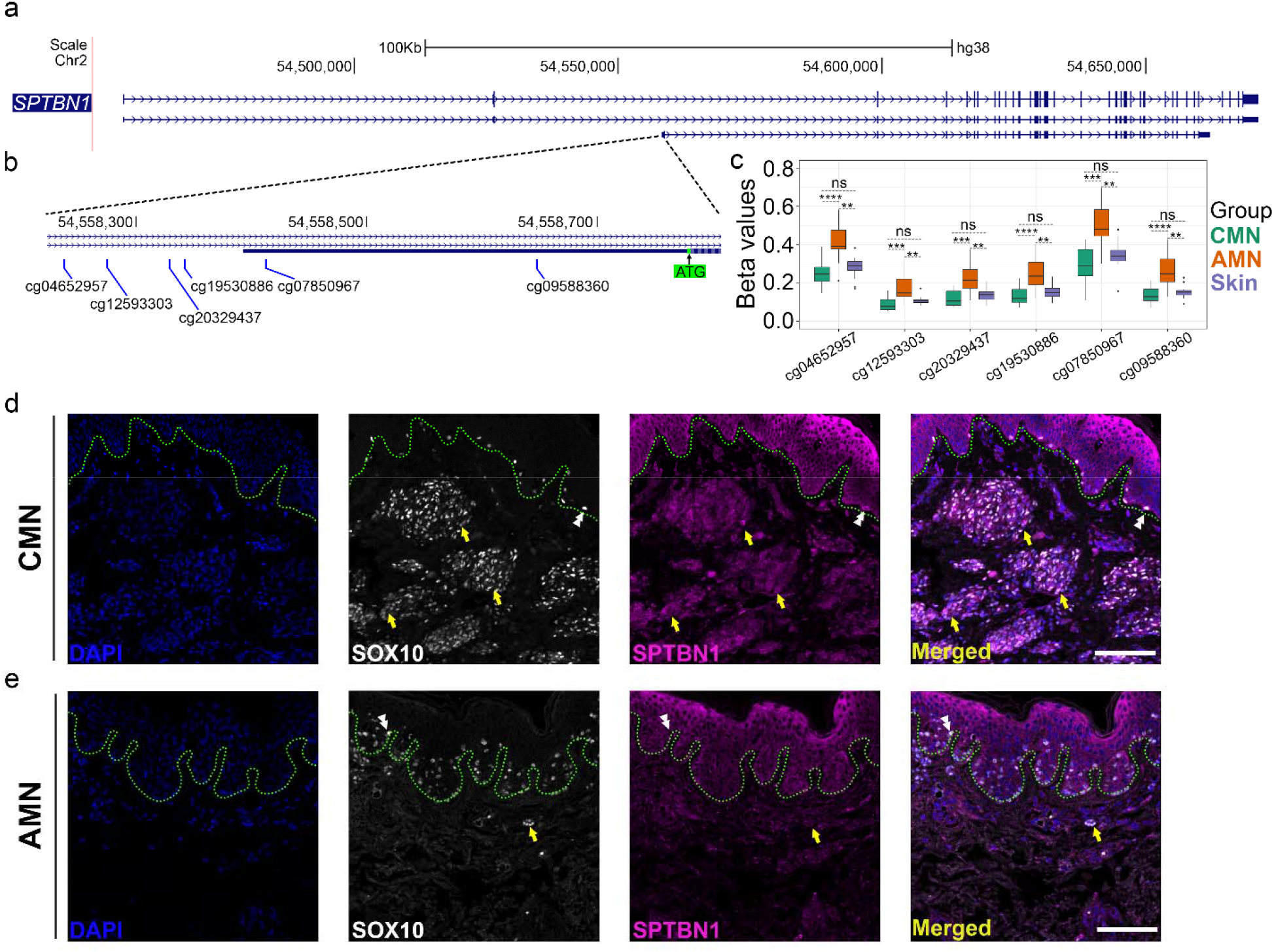
Spectrin Beta, non-erythrocytic 1 (SPTBN1), a general marker of nevomelanocytes. **(a)** DMP genomic positions within the *Spectrin Beta, Non-Erythrocytic 1* (*SPTBN1*) locus. **(c)** Box-plot of the β-values of six DMPs within the *SPTBN1* alternative promoter region. **(d-e)** Representative images of CMN, and AMN stained with antibodies against SPTBN1 (purple) and SOX10 (white), a known melanocyte marker of the superficial skin. Dermal and epidermal melanocytes are indicated by yellow arrows or white double arrowheads, respectively. The dermal-epidermal junction is denoted by dotted lines, nuclei stained by DAPI. Scale bar = 100 µm. In **(c)**, significance was assessed by a Kruskal-Wallis one-way ANOVA. n.s. not significant, **P<0.01, ***P<0.001, ****P<0.0001.

In this study, we investigated the DNA methylation status of pediatric CMN samples compared to a relatively young cohort of AMN and matched adjacent control skin (Muse et al. 2022). By demonstrating fruitful integration of new with existing repository datasets, our study highlighted potentially functional CpGs, like those near the *SPTBN1* gene. Spectrin proteins have been described as playing a role in clustering early perinuclear melanosomes during their maturation (Watabe et al. 2008). Moreover, decreased expression of *SPTBN1* has been associated with tumor progression through the promotion of epithelial-mesenchymal transition in hepatocellular carcinoma, colon cancer, and ovarian cancer (Chen et al. 2020; Mitra et al. 2017; Yokota et al. 2016). Intriguingly, we identified hypo-DMPs near the *RBM24* and *KIF1A* genes (Figure 1g), both of which have been linked with melanoma (Shi 2022; Suresh et al. 2023). Future studies are needed to explore the molecular mechanisms that trigger tumor initiation in CMN and to determine whether the misregulation of *SPTBN1* or other genes plays a driving role in this process.

## DATA AVAILABILITY STATEMENT

Datasets related to this article can be found at [https://doi.org/10.6084/m9.figshare.26005255.v2], hosted at figshare. Methylome data for AMN and control skin are available at GEO under accession GSE188593 (Muse et al. 2022). Methylome data for CMN are available at GEO under accession GSE268769 or released to qualified investigators upon request to the authors. All other data are available in the manuscript or the supplementary materials.

## ACKNOWLEDGMENTS

The authors thank the many patients who agreed to take part in this study. This work was supported by the Horizon Europe Mission Cancer grant 101096667 MELCAYA, the AFM-Téléthon strategic project MoThARD, with additional grants from Nevus Outreach, Inc. (USA), Asociación Española de Afectados por Nevus Gigante Congénito (Spain), Association du Naevus Géant Congénital (France), Institut MarMaRa of Aix-Marseille Université, (France) and Naevus 2000 France-Europe (France).

## AUTHOR CONTRIBUTIONS STATEMENT (CRediT-compliant)

N.M. and H.C.E.: **Conceptualization, Supervision**. D.A., N.M., E.M. and J.G.: **Investigation**. D.A., E.M., and H.C.E.: **Writing – Original Draft**. D.A., N.M., P.R.: **Methodology**. E.M., M.M., and M.P.: **Validation**. D.A., N.M. J.G.: **Formal analysis**. N.D., S.M., I.J. and G.C.: **Resources** (patient enrollment & consent). D. A. and H.C.E.: **Visualization** N.M., A.B. and H.C.E.: **Writing – Reviewing and Editing**; **Funding Acquisition**. H.C.E.: **Project administration**.

## REFERENCES

Bastian BC, Xiong J, Frieden IJ, Williams ML, Chou P, Busam K, et al. Genetic changes in neoplasms arising in congenital melanocytic nevi: differences between nodular proliferations and melanomas. The American Journal of Pathology. 2002;161(4):1163–9

Busam KJ, Shah KN, Gerami P, Sitzman T, Jungbluth AA, Kinsler V. Reduced H3K27me3 Expression is Common in Nodular Melanomas of Childhood Associated with Congenital Melanocytic Nevi but Not in Proliferative Nodules. American Journal of Surgical Pathology. Lippincott Williams and Wilkins; 2017;41(3):396–404

Chen M, Zeng J, Chen S, Li J, Wu H, Dong X, et al. SPTBN1 suppresses the progression of epithelial ovarian cancer via SOCS3-mediated blockade of the JAK/STAT3 signaling pathway. Aging (Albany NY). 2020;12(11):10896–911

Dessars B, De Raeve LE, Morandini R, Lefort A, El Housni H, Ghanem GE, et al. Genotypic and Gene Expression Studies in Congenital Melanocytic Nevi: Insight into Initial Steps of Melanotumorigenesis. Journal of Investigative Dermatology. 2009;129(1):139–47

Macagno N, Etchevers HC, Malissen N, Rome A, Hesse S, Mallet S, et al. Reduced H3K27me3 Expression is Common in Nodular Melanomas of Childhood Associated With Congenital Melanocytic Nevi But Not in Proliferative Nodules. Am J Surg Pathol. 2018;42(5):701–4

Maher NG, Scolyer RA, Colebatch AJ. Biology and genetics of acquired and congenital melanocytic naevi. Pathology. 2023;55(2):169–77

Mitra A, Yan J, Xia X, Zhou S, Chen J, Mishra L, et al. IL6Lmediated inflammatory loop reprograms normal to epithelialLmesenchymal transition+ metastatic cancer stem cells in preneoplastic liver of transforming growth factor beta–deficient β2Lspectrin+/− mice. Hepatology. 2017;65(4):1222

Muse ME, Bergman DT, Salas LA, Tom LN, Tan J-M, Laino A, et al. Genome-Scale DNA Methylation Analysis Identifies Repeat Element Alterations that Modulate the Genomic Stability of Melanocytic Nevi. J Invest Dermatol. 2022;142(7):1893-1902.e7

Pampena R, Kyrgidis A, Lallas A, Moscarella E, Argenziano G, Longo C. A meta-analysis of nevus-associated melanoma: Prevalence and practical implications. J Am Acad Dermatol. 2017;77(5):938-945.e4

Shi D-L. RBM24 in the Post-Transcriptional Regulation of Cancer Progression: Anti-Tumor or Pro-Tumor Activity? Cancers. Multidisciplinary Digital Publishing Institute (MDPI); 2022;14(7) Available from: https://www.ncbi.nlm.nih.gov/pmc/articles/PMC8997389/

Solé-Boldo L, Raddatz G, Schütz S, Mallm J-P, Rippe K, Lonsdorf AS, et al. Single-cell transcriptomes of the human skin reveal age-related loss of fibroblast priming. Commun Biol. Nature Publishing Group; 2020;3(1):1–12

Suresh S, Rabbie R, Garg M, Lumaquin D, Huang T-H, Montal E, et al. Identifying the Transcriptional Drivers of Metastasis Embedded within Localized Melanoma. Cancer Discov. 2023;13(1):194–215

Tirosh I, Izar B, Prakadan SM, Wadsworth MH, Treacy D, Trombetta JJ, et al. Dissecting the multicellular ecosystem of metastatic melanoma by single-cell RNA-seq. Science. 2016;352(6282):189–96

Yang P, Yang Y, Sun P, Tian Y, Gao F, Wang C, et al. βII spectrin (SPTBN1): biological function and clinical potential in cancer and other diseases. International Journal of Biological Sciences. Ivyspring International Publisher; 2021;17(1):32–49

Yokota M, Kojima M, Higuchi Y, Nishizawa Y, Kobayashi A, Ito M, et al. Gene expression profile in the activation of subperitoneal fibroblasts reflects prognosis of patients with colon cancer. International Journal of Cancer. 2016;138(6):1422–31

Zalaudek I, Guelly C, Pellacani G, Hofmann-Wellenhof R, Trajanoski S, Kittler H, et al. The Dermoscopical and Histopathological Patterns of Nevi Correlate with the Frequency of BRAF Mutations. J Invest Dermatol. Elsevier; 2011;131(2):542–5

