## Supplementary for "Deconvoluted methylation profiles discriminate between closely related melanocytic nevi"

### SUPPLEMENTARY FIGURES

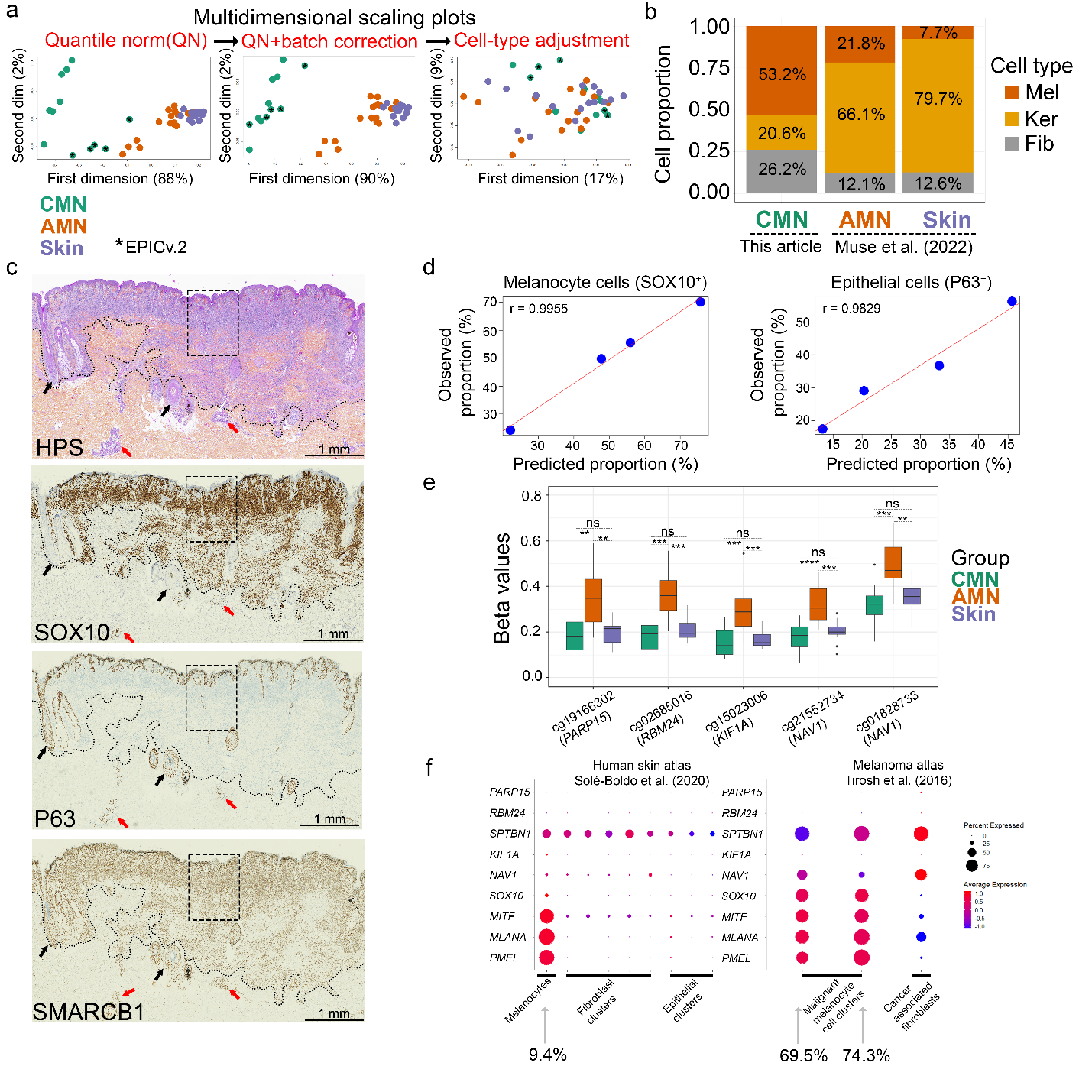

**Supplementary Figure S1.** **(a)** Multidimensional scaling (MDS) plots of DNA methylation profiles from congenital and acquired melanocytic nevi (CMN, AMN), and control (Skin) groups after Quantile normalization, technology batch correction, and cell-type deconvolution. **(b)** Estimated average cellular proportions in CMN, AMN, and control Skin are plotted for melanocytes (Mel), keratinocytes (Ker), and fibroblasts (Fib). **(c)** Representative CMN images. The area shown in Figure 1c is boxed. Hematoxylin-phloxine-saffron (HPS) staining shows high nuclear density (purple) underlying the epidermis. Immunohistochemistry staining with specific markers for (nevo)melanocytes (SOX10^+^), epidermis and epidermal annexes (P63^+^), and a nuclear protein ubiquitously expressed in normal cells (SMARCB1^+^). The melanocytic area (dotted lines), hair follicles (black arrow), and eccrine sweat glands (red arrow) are indicated. **(d)** Pearson coefficient correlation between the observed and predicted proportion of SOX10^+^ melanocyte and P63^+^epidermal cells. **(e)** Box-plot of the beta values of the top hypo-DMPs in CMN vs AMN (see Figure 1f) for Poly(ADP-Ribose) Polymerase Family Member 15 (*PARP15*), RNA Binding Motif Protein 24 (*RBM24*), Kinesin Family Member 1A (*KIF1A*) and Neuron Navigator 1 (*NAV1*). **(f)** Dot plots display the expression of genes associated with the top hypo-DMPs in the CMN versus AMN comparison. The plots highlight normal and malignant melanocytes, fibroblasts, and epithelial cell clusters. Data from atlases were obtained from the Gene Expression Omnibus accessions GSE130973 and GSE72056 (Solé-Boldo et al. 2020; Tirosh et al. 2016). SRY-box transcription factor 10 (*SOX10*), Melanocyte Inducing Transcription Factor (*MITF*), Melan-A (*MLANA*), and Premelanosome Protein (*PMEL*) are gene markers of melanocyte cellular identity. The percentage of cells co-expressing *SOX10* and *SPTBN1* in melanocyte-related clusters is indicated, correspondingly. In **(e)**, significance was assessed by a Kruskal-Wallis one-way ANOVA. n.s. not significant, **P<0.01, ***P<0.001, ****P<0.0001.

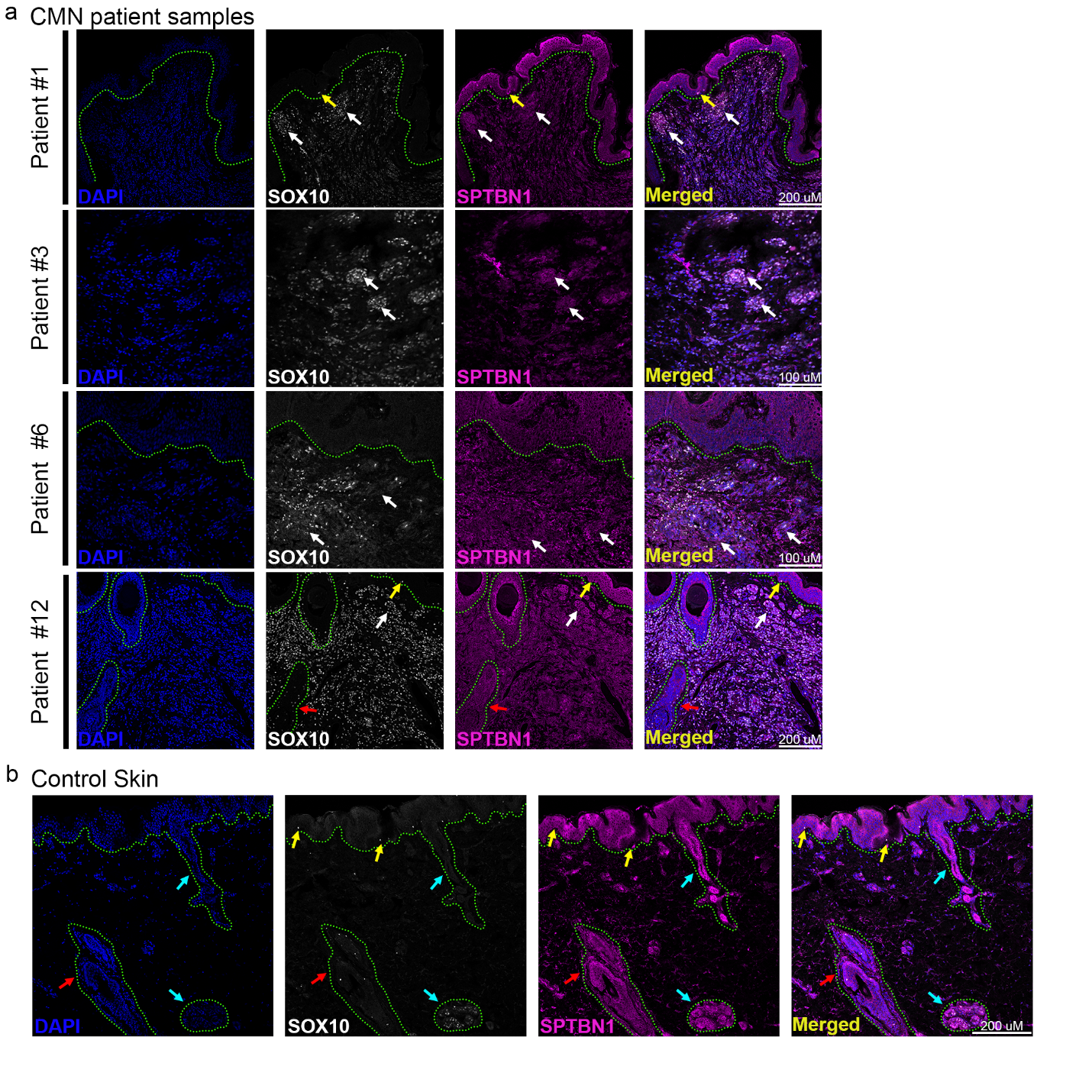

**Supplementary Figure S2.** **(a-b)** Representative images of sections from CMN patients (see Supplementary Table 1) and adjacent unaffected skin from one patient (Control Skin). SPTBN1 and SOX10 proteins were detected through immunofluorescence. The dermal-epidermal junctions (dotted green lines), melanocytic nests (white arrows), normal epithelial melanocytes (yellow arrows), hair follicles (red arrows), and eccrine sweat glands (light-blue arrows) are indicated accordingly.

SUPPLEMENTARY MATERIALS AND METHODS

##### Patients

Samples from twelve patients under the age of 18 years old, affected with small to giant congenital melanocytic nevi (CMN), were taken after independently scheduled resection in reconstructive surgery with written, informed parental consent. They were phenotyped according to (Krengel et al. 2013). To ensure the complete extraction of both superficial (epidermal) and deep (dermal) melanocytes, hematoxylin-phloxine-saffron (HPS) staining was performed on CMN samples and analyzed by an anatomic pathologist. This study was carried out under the aegis of the nationally appointed ethical review board CPP under authorization 214 C03 to H.C.E. and to N.M. by the local ethics committee (APHM, Ref. n°P9X2AQ). Some retrospective consolidated data (samples 8-12) produced during pathology diagnostics were used with a consent waiver granted for this work.

##### Methylation array hybridization

DNA was extracted from flash-frozen biopsies or paraffin blocks using standard salting-out techniques. DNA concentrations were determined by the Qubit® (ThermoFisher) fluorometric assay. Bisulfite treatment of 500 ng DNA to convert unmethylated cytosine nucleotides to uracil was carried out using the EpiJET Bisulfite Conversion Kit (ThermoFisher Scientific) and eluted in 10 µL. After normalization and denaturation, DNAs were hybridized to an Infinium Methylation EPIC.v1 or EPIC.v2 BeadChip arrays (Illumina) targeting over 850,000 CpG sites encompassing RefSeq genes, CpG islands, enhancers, ENCODE open chromatin and transcription factor binding sites, and miRNA promoters, according to manufacturer’s instructions, by IntegraGen (Evry, France) or Eurofins Genomics SAS (Nantes, France), respectively. After overnight hybridization with rocking at 48ºC, the array was washed, stained, the array surface sealed, then scanned using an Illumina iSCAN platform, according to the manufacturer’s instructions.

##### DNA methylation analysis

###### Data processing

Preprocessing and quality control of the sample intensity data (IDAT) files were performed using the R package Minfi package (v.1.48.0) (Aryee et al. 2014). To account for inter-laboratories batch effects, the samples were normalized together using the preprocessQuantile function from the same package. Poor-quality probes (p-value <0.01) and probes associated with Single Nucleotide Polymorphism (SNPs) or sex-chromosomes were identified and excluded. Our data set included two types of arrays: MethylationEPIC v1 (>850,000 CpG probes) and MethylationEPIC v2 (>935,000 CpG probes) (see Supplementary Table S1). To combine arrays, beta values from the MethylationEPIC v2 array were converted to MethylationEPIC v1 array format using the liftover function from the Sesame package (v.1.20.0) (Zhou et al. 2018). Potential batch effects resulting from the two types of microarrays (MethylationEPIC v1 and MethylationEPIC v2) were corrected using the ComBat function from the *sva* package in R (Johnson et al. 2007; Leek et al. 2012) (Supplementary Fig. 1a).

Cell-type deconvolution

Relative proportions of component cell types included in our samples were estimated using the Epidish function (Teschendorff et al. 2017), using the reference library for DNA methylation-based cell-type deconvolution of skin biopsies implemented by (Muse et al. 2022). This library includes epidermal cells (keratinocytes), dermal fibroblasts, and melanocytes among others. After filtering, normalization, batch correction, and cell deconvolution steps, a total of 698,939 CpG probes for each sample were used for downstream analysis.

###### Differentially Methylated Positions

Differentially Methylated Positions (DMPs) at unique CpGs were identified through mixed linear regression models using the *limma* package from R (Ritchie et al. 2015). In brief, in a pairwise comparison for the CMN, AMN, and unaffected Skin groups, the fold changes and standard errors of each CpG were estimated by fitting a linear model followed by Empirical Bayes smoothing and using the logit-transformed beta values (m-values) as input. This analysis was performed with the Limma 3.58.1 package. DMPs with P-value adjusted <0.05 (False Discovery Rate, FDR), and a size effect (log2FC) |≥0.5| were considered to be hyper- or hypo-methylated probes.

##### Single cell RNA-seq analysis

Seurat R package (version 5) (Hao et al. 2024) was used to analyze the single-cell RNA-seq datasets. Skin atlases were downloaded from the Gene Expression Omnibus (accessions GSE130973 and GSE72056) (Solé-Boldo et al. 2020; Tirosh et al. 2016). Gene expression levels were plotted using the Dotplot function from the same package across various cell clusters from both normal and melanoma human skin atlases. These clusters included melanocytes (85 cells), fibroblasts (6,131 cells), and epithelial cells (2,822 cells) from normal skin, as well as malignant melanocytes (1,758 cells) and cancer-associated fibroblasts (61 cells) from melanoma samples. Cell clustering was previously annotated and was not modified (Solé-Boldo et al. 2020; Tirosh et al. 2016). Details of these atlases can be found at [ <https://doi.org/10.6084/m9.figshare.26005255.v2> ], hosted at figshare.

##### Immunofluorescence and imaging

After fixation in formalin, skin tissues were embedded in paraffin and sectioned at 3.5 or 5 µm before immunostaining processing. Briefly, tissue sections from seven CMN patients were deparaffinized in a graded series of xylenes then ethanols to water, post-fixed in 4% buffered paraformaldehyde (PFA) and washed in phosphate-buffered saline (PBS). Epitopes were exposed after a heat-inactivation antigen retrieval step according to primary antibody recommendations. Pigments were bleached and endogenous peroxidases blocked using 0.3% hydrogen peroxide in water and aldehyde quenching was performed using PBS-glycine-ammonium acetate. Next, tissue was blocked in PBST (PBS + 0.1% Tween) + 2% normal bovine serum before incubating in 1:100 each dilutions of mouse monoclonal anti-SPTBN1 primary antibody (clone 1E1D1, Proteintech, cat. 67978-1-Ig) or rabbit monoclonal anti-SOX10 primary antibody (clone EP268, GeneTex, cat. GTX03312). Secondary detection was performed with species-specific antibodies conjugated to Alexa Fluor-555 (1:250, Invitrogen) or Alexa Fluor-647 (1:250, Invitrogen), and samples were counterstained with 4′6-diamidino-2-phenylindole (DAPI). Images were acquired on a Zeiss LSM 800 Confocal using the ZEISS ZEN 3.7 Base version 3.97.23089.111. Immunohistochemistry was performed using an automated system (Benchmark ULTRA; Ventana Medical Systems Inc., Tucson, AZ) for P63 (clone: 4A4, Ventana), SOX10 (clone SP267, Ventana) and SMARCB1(INI-1) (clone 25, Bio SB, Santa Barbara, CA) using standard procedures. All images shown in this article can be found at [ <https://doi.org/10.6084/m9.figshare.26005255.v2> ], hosted at figshare.

**Semi-automatic cell number determination**

Image segmentation was performed using “ilastik: interactive machine learning for (bio)image analysis”, v1.4.0.post1 (Berg et al. 2019). In brief, cell scoring was performed using immunohistochemistry images following the nuclear density counting workflow from ilastik. The percentages of melanocytes (nuclear SOX10^+^) and keratinocytes (P63^+^) were calculated as a proportion of the total cellular number (SMARCB1^+^).

**Statistical analysis**

Statistical analysis was performed by means of two-tailed Mann-Whitney T-test or Kruskal-Wallis one-way ANOVA tests using GraphPad Prism version 10.2. for Windows (GraphPad Software, La Jolla, CA).

### SUPPLEMENTARY REFERENCES

Aryee MJ, Jaffe AE, Corrada-Bravo H, Ladd-Acosta C, Feinberg AP, Hansen KD, et al. Minfi: a flexible and comprehensive Bioconductor package for the analysis of Infinium DNA methylation microarrays. Bioinformatics. 2014;30(10):1363–9

Berg S, Kutra D, Kroeger T, Straehle CN, Kausler BX, Haubold C, et al. ilastik: interactive machine learning for (bio)image analysis. Nat Methods. Nature Publishing Group; 2019;16(12):1226–32

Hao Y, Stuart T, Kowalski MH, Choudhary S, Hoffman P, Hartman A, et al. Dictionary learning for integrative, multimodal and scalable single-cell analysis. Nat Biotechnol. 2024;42(2):293–304

Johnson WE, Li C, Rabinovic A. Adjusting batch effects in microarray expression data using empirical Bayes methods. Biostatistics. 2007;8(1):118–27

Krengel S, Scope A, Dusza SW, Vonthein R, Marghoob AA. New recommendations for the categorization of cutaneous features of congenital melanocytic nevi. J Am Acad Dermatol. 2013;68(3):441–51

Leek JT, Johnson WE, Parker HS, Jaffe AE, Storey JD. The sva package for removing batch effects and other unwanted variation in high-throughput experiments. Bioinformatics. 2012;28(6):882–3

Muse ME, Bergman DT, Salas LA, Tom LN, Tan J-M, Laino A, et al. Genome-Scale DNA Methylation Analysis Identifies Repeat Element Alterations that Modulate the Genomic Stability of Melanocytic Nevi. J Invest Dermatol. 2022;142(7):1893-1902.e7

Ritchie ME, Phipson B, Wu D, Hu Y, Law CW, Shi W, et al. limma powers differential expression analyses for RNA-sequencing and microarray studies. Nucleic Acids Research. 2015;43(7):e47

Solé-Boldo L, Raddatz G, Schütz S, Mallm J-P, Rippe K, Lonsdorf AS, et al. Single-cell transcriptomes of the human skin reveal age-related loss of fibroblast priming. Commun Biol. Nature Publishing Group; 2020;3(1):1–12

Teschendorff AE, Breeze CE, Zheng SC, Beck S. A comparison of reference-based algorithms for correcting cell-type heterogeneity in Epigenome-Wide Association Studies. BMC Bioinformatics. 2017;18(1):105

Tirosh I, Izar B, Prakadan SM, Wadsworth MH, Treacy D, Trombetta JJ, et al. Dissecting the multicellular ecosystem of metastatic melanoma by single-cell RNA-seq. Science. 2016;352(6282):189–96

Zhou W, Triche TJ Jr, Laird PW, Shen H. SeSAMe: reducing artifactual detection of DNA methylation by Infinium BeadChips in genomic deletions. Nucleic Acids Research. 2018;46(20):e123
