## Supplementary figures and images for "Deconvoluted methylation profiles discriminate between closely related melanocytic nevi"

### Supplementary Fig. S1

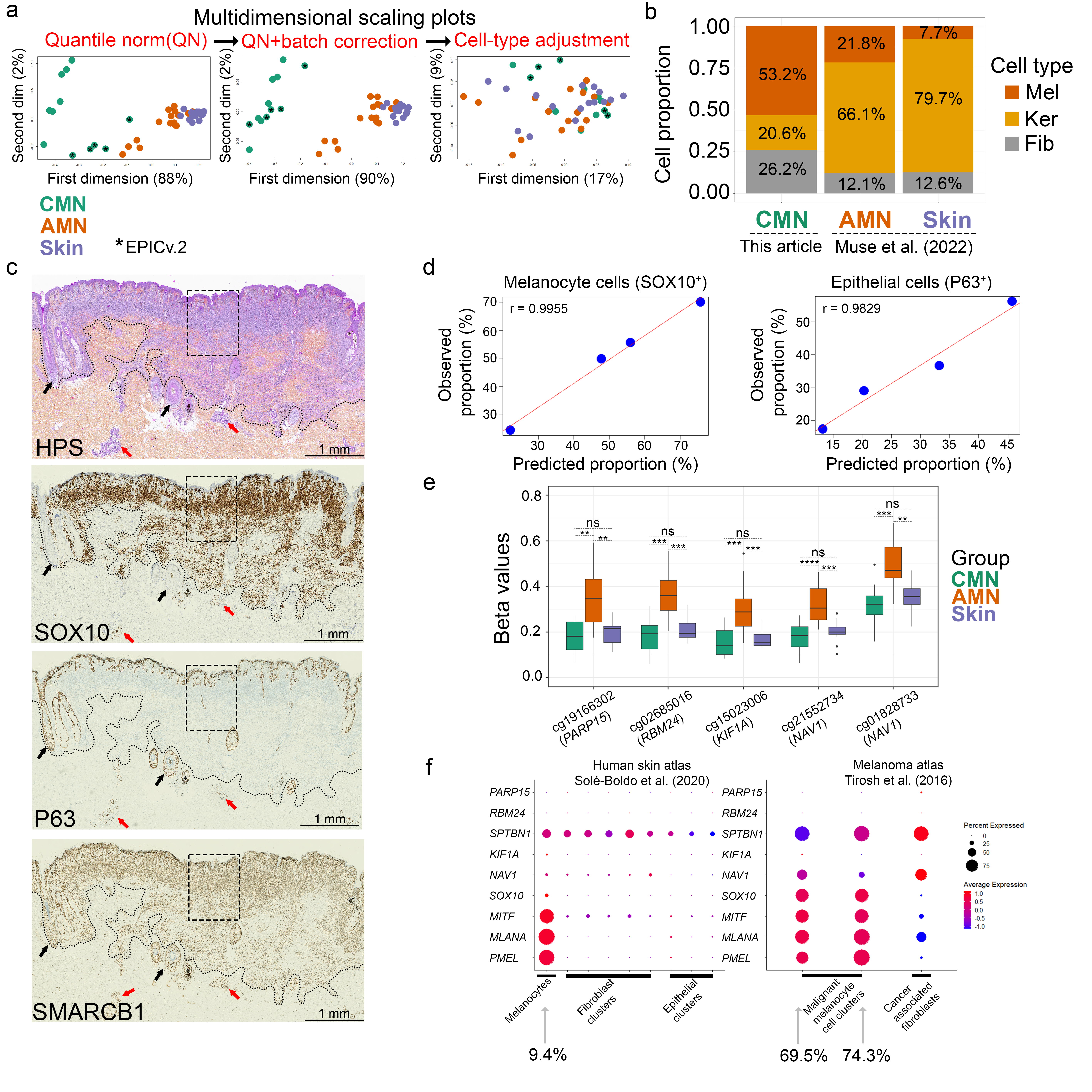

### Supplementary Fig. S2

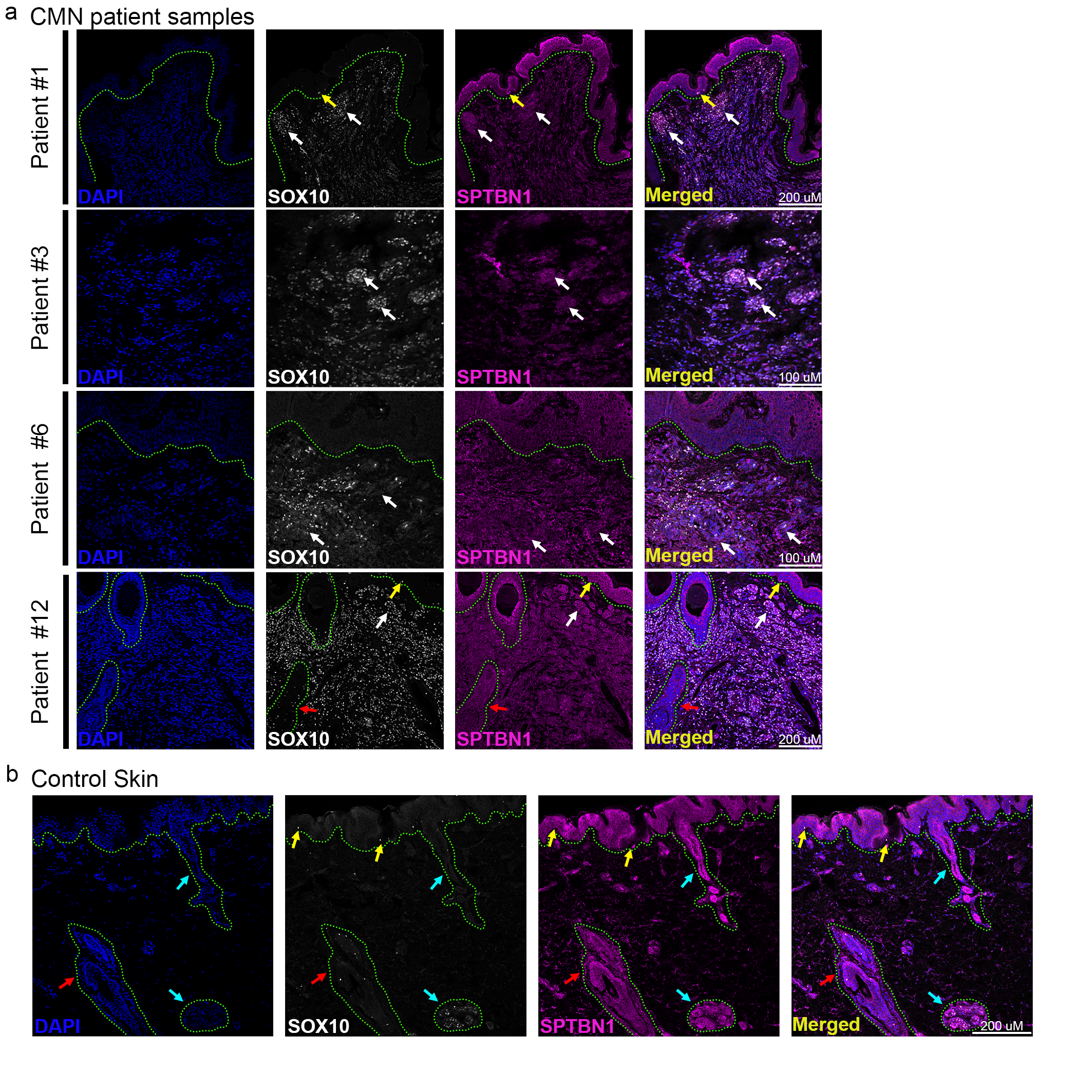
